# A long shelf-life melon created via CRISPR/Cas9 RNP-based *in planta* genome editing

**DOI:** 10.1101/2024.09.16.613271

**Authors:** Kentaro Sasaki, Naozumi Mimida, Satoko Nonaka, Hiroshi Ezura, Ryozo Imai

**Affiliations:** Institute of Agrobiological Sciences, National Agriculture and Food Research Organization (NARO), Tsukuba, Japan; Sanatech Life Science Co. Ltd., Minato-ku, Tokyo, Japan; Tskuba Plant Innovation Research Center, University of Tsukuba, Tsukuba, Japan; Faculty of Life and Environmental Sciences, University of Tsukuba, Tsukuba, Japan

**Author notes:** **Corresponding author:** Ryozo Imai *E-mail address:.

**Keywords:** CRISPR/Cas9, ethylene, genome editing, melon, shoot apical meristem, shelf-life

## Abstract

We developed a non-culture and DNA-free genome editing technique for melon using in plant Particle Bombardment (iPB-RNP). The method was employed to create CmACO1 mutants. One of the mutant lines exhibited an extended shelf life due to ethylene deficiency during fruit ripening. As no cell culture step is involved, the iPB-RNP method is expected to overcome limitations associated with conventional genome editing, such as genotype dependency and somatic variations. Thus, this methodology holds significant potential for application in commercial melon breeding and across a wide range of Cucurbitaceae species.

---

**M**elon (*Cucumis melo* L.) is a widely consumed fruit globally. Although conventional breeding has enhanced the quality and productivity of melon varieties, there is a pressing need for rapid and diverse genetic improvements. Genome editing is anticipated to fulfill these demands. Most current genome editing protocols depend on genetic transformation and tissue culture, yet melon exhibits low transformation efficiency due to its limited regeneration capabilities and the occurrence of escapes (Shirazi Parsa et al., 2023). Additionally, melon is prone to ploidy changes during tissue culture, altering morphogenesis in regenerated plants (Nonaka et al., 2023; Nuñez-Palenius et al., 2008). Therefore, an efficient and robust system for melon genetic enhancement through genome editing is required.

In a prior study, we developed a transgene-free genome editing approach, in planta particle bombardment-ribonucleoprotein (iPB-RNP), applied in wheat and barley (Kumagai et al., 2022; Tezuka et al., 2024). This technique employs the CRISPR/Cas9 ribonucleoprotein, introduced directly into the shoot apical meristem (SAM) to specifically target subepidermal L2 cells, potential germline cells (Goldberg et al., 1993). Since it bypasses cell culture and regeneration processes, iPB-RNP could be applicable to recalcitrant crop species, including melon.

To assess the feasibility of genome editing in melon via iPB-RNP, we first evaluated the efficiency of gold particle delivery into the meristematic cells. Following 20 hours of imbibition, we removed half of the cotyledon from germinated seeds (cv. “Harukei-3”) to expose the SAM, and positioned the remaining seeds on an agar plate with the exposed SAM facing upwards (Figure 1a). Gold particles coated with a GFP-expressing vector were bombarded towards the SAMs on the plate (Figure 1a). After 16 hours in darkness, GFP signals were observed on the SAM surface in almost all bombarded plants, unlike the controls (Figure 1a and S1). These results confirmed the proper preparation of SAMs and the suitability of the bombardment conditions for melon iPB.

**Figure 1.**
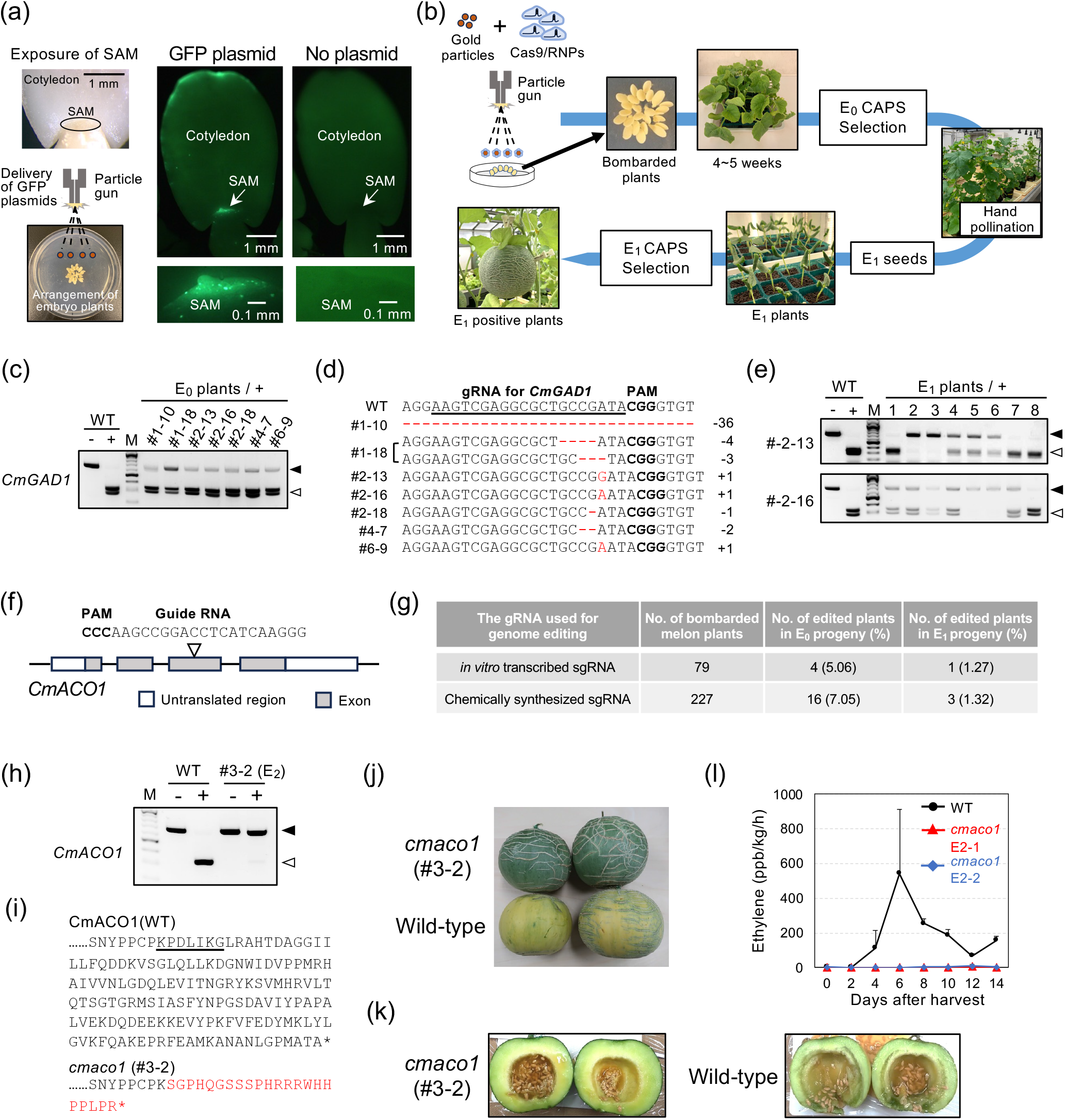
*In planta* RNP-mediated genome editing in melon. (a) Microprojectile-mediated GFP gene transfer to melon SAMs. (b) Overview of the iPB-RNP method for genome editing in melon. (c) CAPS analysis of positive E_0_ plants carrying mutations at the targeted *CmGAD1* locus. Symbols “–” and “+” represent digestion without and with Cas9 RNPs, respectively. Black and white triangles indicate undigested and digested bands following Cas9 RNP treatment, respectively. (d) Alignment of CRISPR/Cas9 target sequences from positive E_0_ plants with that of the wild-type, with insertions and deletions highlighted in red letters. (e) Gene-specific CAPS analysis of E_1_ plants derived from two E_0_ plants (#2-13 and #2-16), using the same symbols as in (c). (f) Schematic of the *CmACO1* gene with gRNA design. (g) Summary of genome editing experiments targeting *CmACO1*. (h) Gene-specific CAPS analysis of the *cmaco1* homozygous mutant line (#3-2, E_2_ generation), using symbols consistent with (c). (i) Amino acid sequence comparison of CmACO1 in wild-type and the *cmaco1*, with target sites for CRISPR/Cas9 underlined and sequence changes in red letters. An asterisk denotes the stop codon. (j and k) Appearance (j) and longitudinal section (k) of fruit from wild-type and the *cmaco1* post-harvest. (l) Ethylene production in fruits from wild-type and *cmaco1* (individual plants E2-1 and E2-2 in the E_2_ generation) measured 40 days after pollination. Data are presented as means ± SE (n = 3).

To implement the iPB-RNP method in melon, we targeted *CmGAD1*, a glutamate decarboxylase gene (You et al., 2024). Preassembled SpCas9 RNP targeting *CmGAD1* (Figure S2) was coated onto gold particles and delivered to the SAM of germinated seeds. The plants grown from the bombarded seeds were analyzed after 4-5 weeks, and their 6th leaf was subjected to cleaved amplified polymorphic sequences (CAPS) analysis for mutation detection (Figure 1b). Seven out of 153 initially bombarded plants showed undigested bands in the CAPS assay (Figure 1c and Table S1), with subsequent sequencing confirming mutations at the target site (Figure 1d). Since the E0 plants exhibit chimeric mutations, E1 plants were analyzed to confirm fixed mutations. The CAPS assay detected a mutant allele of *CmGAD1* in two out of the seven E0 progenitors (Figure 1e and Table S1). Among these, #2-13_2, #2-13_3, #2-16_5, and #2-16_6 were homozygous mutants for *CmGAD1* (Figure 1e), and sequence analysis showed that mutations in the E1 plants were consistent with those detected in the E0 generation (Figure 1d and S3). The efficiency of iPB-RNP editing for *CmGAD1* was 1.31% (two out of 153 bombarded plants), comparable to efficiencies observed in wheat and barley (Kumagai et al., 2022; Tezuka et al., 2024).

Having established the iPB-RNP in melon, we then applied this method to extend the shelf-life of melon, a desirable trait. Since the gaseous phytohormone ethylene controls fruit maturation, our target was CmACO1, an ethylene biosynthesis gene (Nonaka et al., 2023). We designed a guide RNA for *CmACO1* (Figure 1f) and synthesized it either enzymatically or chemically (Figure 1g). Cas9 RNPs targeting *CmACO1* were delivered into SAMs by iPB-RNP. Ultimately, we obtained one and three genome-edited lines, corresponding to 1.27% and 1.32% of the total, with enzymatically and chemically synthesized gRNAs, respectively (Figure 1g and S4).

Given the role of CmACO1 in fruit ripening, we compared the fruit phenotype between the wild-type and the #3-2 homozygous mutant (Figure 1h). The #3-2 line harbors a one-base insertion within the *CmACO1* gene, suggesting a loss of CmACO1 function (Figure 1i and S4f). The fruit size and shape of the *cmaco1* mutant were comparable to those of the wild-type. Ten days after harvest, the pericarp color changed from green to yellow in the wild-type fruit but remained green in the *cmaco1* mutant (Figure 1j). Cross-sectional analysis of the fruit after harvest revealed a mushy texture in the wild-type, which was absent in the *cmaco1* mutant (Figure 1k). To confirm that the delay in fruit ripening was caused by reduced ethylene production in the *cmaco1* mutant, ethylene levels were measured post-harvest. Wild-type fruit exhibited a significant ethylene burst, peaking 6 days after harvest, whereas the *cmaco1* mutant consistently produced significantly lower ethylene levels up to 14 days after harvest (Figure 1l). In a previous study, a *cmaco1* mutant generated by conventional Agrobacterium-mediated transformation displayed smaller and flattened fruit due to somatic mutations that induced tetraploidy (Nonaka et al., 2023). Our findings demonstrate that the iPB-RNP method effectively circumvents complications associated with cell cultures, ultimately enabling the creation of melons with an extended shelf life without inducing morphological or growth defects.

In summary, we developed a non-culture and DNA-free editing technique for melon using iPB-RNP. The efficiency for obtaining positive plants in the E1 generation reached 1.27% and 1.32% for *CmACO1*, and 1.31% for *CmGAD1* (Figure 1g and Table S1). This editing efficiency in melon is comparable with those in wheat and barley (Kumagai et al., 2022; Tezuka et al., 2024). As no cell culture step is involved, the iPB-RNP method is expected to overcome limitations associated with conventional genome editing, such as genotype dependency and somatic variations (Nuñez-Palenius et al., 2008). Thus, this methodology holds significant potential for application in commercial melon breeding and across a wide range of Cucurbitaceae species.

## 1. Acknowledgements

This work was supported by the Cross-ministerial Strategic Innovation Promotion Program (SIP), “Technologies for Smart Bio-industry and Agriculture”, with funding provided by the Bio-oriented Technology Research Advancement Institution to H.E. and R.I.

## 2. Author contributions

H.E., and R.I. designed research; K.S., N.M., and S.N. performed research; K.S., N.M., H.E., and R.I. analyzed data; and K.S. and R.I. wrote the paper.

## Notes

### Competing Interest Statement

The authors have declared no competing interest.

### Summary of Updates

The manuscript has been formatted to improve the visibility.

## References

Goldberg, R.B., Beals, T.P., and Sanders, P.M. (1993) Anther development: Basic principles and practical applications. Plant Cell, 5, 1217–1229.

Kumagai, Y., Liu, Y., Hamada, H., Luo, W., Zhu, J., Kuroki, M., et al. (2022) Introduction of a second “Green Revolution” mutation into wheat via in planta CRISPR/Cas9 delivery. Plant Physiol., 188, 1838–1842.

Nonaka, S., Ito, M., and Ezura, H. (2023) Targeted modification of CmACO1 by CRISPR/Cas9 extends the shelf-life of Cucumis melo var. reticulatus melon. Front. Genome Ed., 5, 1–17.

Nuñez-Palenius, H.G., Gomez-Lim, M., Ochoa-Alejo, N., Grumet, R., Lester, G., and Cantliffe, D.J. (2008) Melon fruits: Genetic diversity, physiology, and biotechnology features. Crit. Rev. Biotechnol., 28, 13–55.

Shirazi Parsa, H., Sabet, M.S., Moieni, A., Shojaeiyan, A., Dogimont, C., Boualem, A., and Bendahmane, A. (2023) CRISPR/Cas9-Mediated Cytosine Base Editing Using an Improved Transformation Procedure in Melon (Cucumis melo L.). Int. J. Mol. Sci., 24, 11189.

Tezuka, D., Cho, H., Onodera, H., Linghu, Q., Chijimatsu, T., Hata, M., and Imai, R. (2024) Redirecting barley breeding for grass production through genome editing of Photoperiod-H1. Plant Physiol., 195, 287–290.

You, W., Zhang, J., Ru, X., Xu, F., Wu, Z., Jin, P., et al. (2024) Cm-CML11 interacts with CmCAMTA5 to enhance gamma-aminobutyric acid (GABA) accumulation by regulating GABA shunt in fresh-cut cantaloupe. Plant Physiol. Biochem., 206, 108217.

