## Supplemental Figs and Tables for "A long shelf-life melon created via CRISPR/Cas9 RNP-based *in planta* genome editing"

(a)

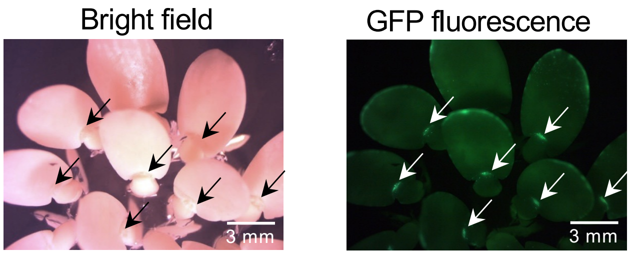


(b)

| Expt. | No. of bombarded plants | No. of plants carrying GFP within SAM (%) |
|---|---|---|
| 1 | 14 | 13 (92.9) |
| 2 | 14 | 14 (100) |

**Supporting Figure S1** Microprojectile-mediated GFP gene transfer to melon SAMs. (a) Bright field and fluorescence images of plants 16 hours post-bombardment, with SAMs highlighted by arrows. (b) Microprojectile-mediated delivery efficiency of the GFP plasmid in melon.

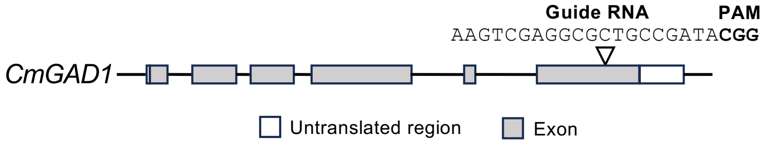


**Supporting Figure S2** *CmGAD1* gene with gRNA design. This schematic provides an overview of the gene structure and the regions targeted by the guide RNA (gRNA).

**gRNA for *CmGAD1*** **PAM**

| | | |
|---|---|---|
| WT | AGGAAGTCGAGGCGCTGCCGATA**CGG**GTGT | |
| #2-13 ($E_1$) | AGGAAGTCGAGGCGCTGCCGGATA**CGG**GTGT | +1 |
| #2-16 ($E_1$) | AGGAAGTCGAGGCGCTGCCGAATA**CGG**GTGT | +1 |

**Supporting Figure S3** Sequence evaluation of *cmgad1* mutant lines in the $E_1$ generation. The CRISPR/Cas9 target sequence of the positive $E_1$ plants (#2-13 and #2-16) is aligned with that of the wild-type, with insertions highlighted in red letters.

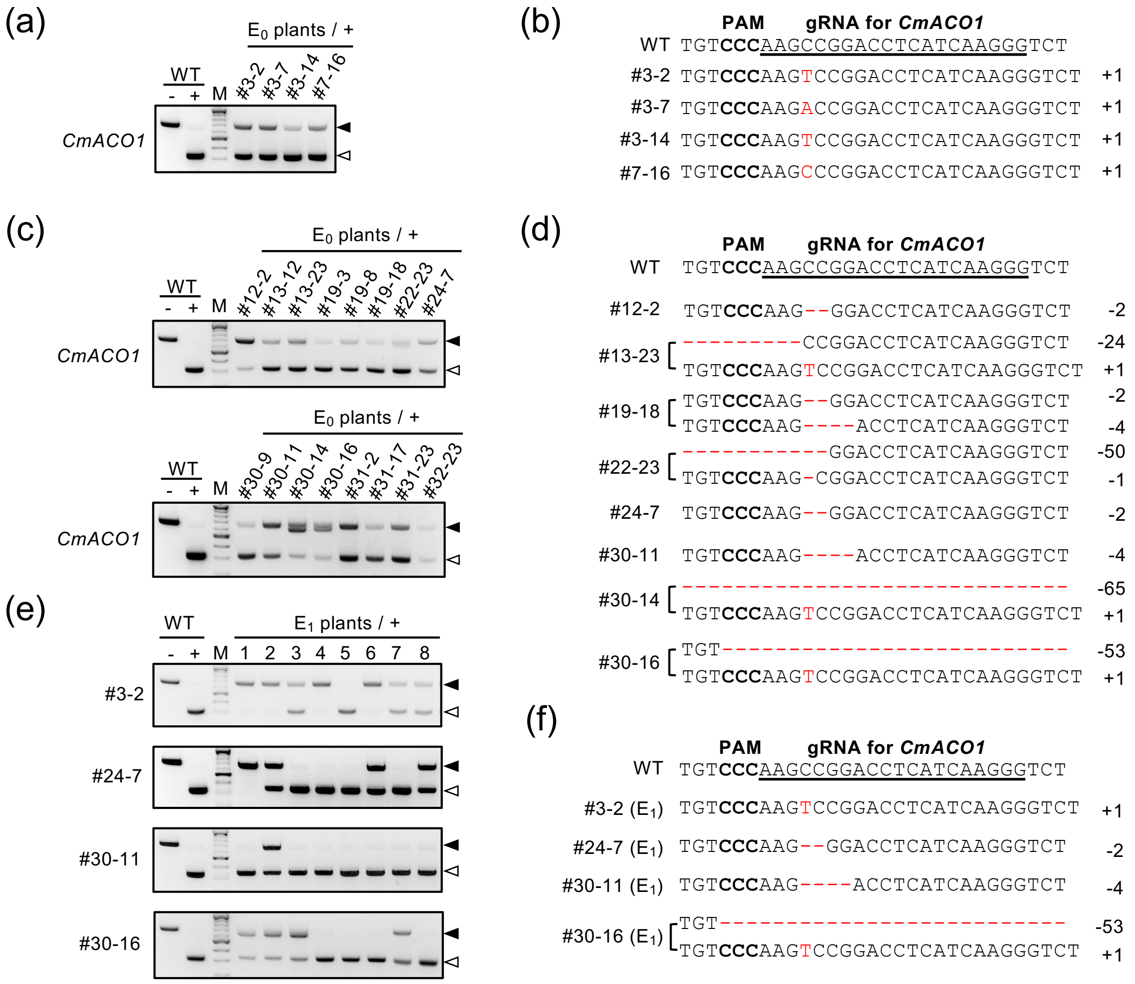


**Supporting Figure S4** Introduction of a mutation in *CmACO1*. (a) CAPS analysis of positive $E_0$ plants created using *in vitro*-transcribed sgRNA. Symbols "–" and "+" denote digestion without and with Cas9 RNPs, respectively. Black and white triangles represent undigested and digested bands after Cas9 RNP treatment, respectively. (b) The CRISPR/Cas9 target sequences of positive $E_0$ plants in (a) are aligned with the wild-type sequence. Insertions are highlighted in red letters. (c) CAPS analysis of positive $E_0$ plants generated using chemically synthesized sgRNA. The symbols have the same meaning as in (a). (d) The CRISPR/Cas9 target sequences of some positive $E_0$ plants in (c) are aligned with the wild-type sequence. Insertions and deletions are indicated in red letters. (e) Gene-specific CAPS analysis of $E_1$ plants derived from four positive $E_0$ plants (#3-2, #24-7, #30-11, and #30-16). The symbols are the same as in (a). (f) The CRISPR/Cas9 target sequence of positive $E_1$ plants (#3-2, #24-7, #30-11, and #30-16) is aligned with that of the wild-type. The symbols are the same as in (d).

**Supporting Table S1** Summary of the genome editing experiment targeting *CmGAD1*. This table presents the editing efficiency calculated based on the number of bombarded plants.

| Target gene | No. of bombarded melon plants | No. of edited plants in $E_0$ progeny (%) | No. of edited plants in $E_1$ progeny (%) |
|---|---|---|---|
| *CmGAD1* | 153 | 7 (4.58) | 2 (1.31) |

**Supporting Table S2** Primers used in this study.

| Primer name | Primer sequence | Note | Purpose |
|---|---|---|---|
| CmGAD1-2Fw | GGGTACAAAAACGTGATGGAAAACTG | For $E_0$ and $E_1$ screening | To amplify the *CmGAD1* target region |
| CmGAD1-2Rv | TGGTGGAAAAAGGGTGTGTATTGTG | | |
| CmACO1-1Fw | TGGGAAAGCACCTTTTTCTTACGCCA | For $E_0$ and $E_1$ screening | To amplify the *CmACO1* target region |
| CmACO1-1Rv | ATCGACATTCGCCCAGTTCC | | |
| CmGAD1_gRNA_Fw | TAATACGACTCACTATAGAAGTCGAGGC GCTGCCG | CmGAD1 | Template DNA amplification to synthesize gRNA for CAPS analysis |
| CmGAD1_gRNA_Rv | TTCTAGCTCTAAAACTATCGGCAGCGCC TCGACTT | | |
| CmACO1_gRNA_Fw | TAATACGACTCACTATAGCCCTTGATGA GGTCCGG | CmACO1 | Template DNA amplification to synthesize gRNA for genome editing and CAPS analysis |
| CmACO1_gRNA_Rv | TTCTAGCTCTAAAACAAGCCGGACCTCA TCAAGGG | | |

**Supporting Table S3** Guide RNA target sites.

| Target gene | Guide RNA sequence | PAM |
|---|---|---|
| *CmGAD1* | AAGTCGAGGCGCTGCCGATA | CGG |
| *CmACO1* | CCCTTGATGAGGTCCGGCTT | GGG |

**Supporting Table S4** Chemically synthesized guide RNA used in this study

| Name | Sequence | Note |
|---|---|---|
| CmGAD1 gRNA | AAGUCGAGGCGCUGCCGAUAGUUUUAGAGCUAUGCUGUUUUG | crRNA |
| tracrRNA | AAACAGCAUAGCAAGUUAAAAUAAGGCUAGUCCGUUAUCAACUU GAAAAAGUGGCACOGAGUCGGUGCU | tracrRNA |
| CmACO1 gRNA | CCCUUGAUGAGGUCCGGCUUGUUUUAGAGCUAGAAAUAGCAAG UUAAAAUAAGGCUAGUCCGUUAUCAACUUGAAAAAGUGGCACCG AGUCGGUGCUUUU | single gRNA |
